# Quantification of Phytohormones in Plants – Optimized Extraction, Separation and Detection

**DOI:** 10.64898/2026.03.17.712349

**Authors:** Vera Wewer, Nadine Dyballa-Rukes, Sabine Metzger

## Abstract

Phytohormones are key players in the regulation of plant development and metabolism. The different phytohormone classes comprise numerous chemically very diverse compounds, which are often present at very low concentrations. The chemical properties of phytohormones range from acidic to basic and from polar to non-polar. Furthermore, concentration varies strongly among different phytohormones, between plant species, tissues and developmental stages. Challenges often arise when only small amounts of plant material are available and when plant species are investigated in which the phytohormone profile has not yet been characterized. To establish a method for comprehensive phytohormone analysis we addressed these challenges by choosing and optimizing a suitable extraction method followed by optimized HPLC separation. We compared the most widely-used mass spectrometric detection methods, multiple reaction monitoring (MRM) on a triple quad instrument with high-resolution mass spectrometry (HRMS) on a Q-TOF instrument, and discuss the advantages of both methods and their limitations.

- We compared various methods described in literature for the extraction of six phytohormone classes by liquid-liquid extraction and solid phase extraction purification and describe our optimizations to the selected method.
- We optimized HPLC separation for 50 different phytohormones.
- We evaluated the application of MRM and HRMS detection strategies.

## Background

Among the pyhtohormones we differentiate between auxins, cytokinins, gibberellins (GA), abscisic acid (ABA), salicylic acid (SA), jasmonates, brassinosteroids, strigolactones and ethylene. The classification of phytohormones gives a first indication that these classes present a great chemical diversity (acidic versus basic, hydrophilic versus hydrophobic, volatile versus non-volatile). In addition, there is high diversity of physicochemical properties even within the same phytohormone class. Many classes include biosynthetic precursors, conjugated forms and isomers, in addition to the bioactive forms. Conjugated phytohormones may be used for transport or storage to quickly release the bioactive compound when required. For example, within the auxins conjugation of indole-3-acetic acid (IAA) with amino acids leads to higher polarity. The same is true for cytokinins, where e.g. zeatin can be conjugated to one or more sugar moieties. Changing physicochemical properties by conjugation has an effect on extraction efficiencies, separation and detection.

To obtain a comprehensive overview of the phytohormone status of a plant, the analysis of not only the bioactive forms but also related molecules of a given phytohormone class will yield important additional information. To achieve this, the selection of a suitable strategy for extraction of all phytohormones from plant tissues, HPLC separation and detection is a challenging task, which we will address in this article.

In early years, phytohormones were analyzed by methods such as immunoassays (reviewed in [1]) which can quantify phytohormones present at higher concentrations. More recent methods show a higher sensitivity and usually comprise chromatographic separation (either by gas chromatography (GC) or LC) coupled to mass spectrometric (MS) detection. With regard to the chromatographic separation, GC is successfully used for volatile phytohormones such as IAA and GA, and the gaseous phytohormone ethylene [2]. Due to the high sensitivity and broad applicability to the chemically diverse phytohormones, LC-MS has become the most widely-used method in current phytohormone research [3].

A multitude of different methods are employed by researchers for sample extraction, optional purification and/or enrichment and HPLC separation (recently reviewed in [4]). The methods have to be adapted to the phytohormones of interest and the plant material. Purification and enrichment strategies are often employed when the focus of the study is on a limited amount of phytohormones, in contrast to other studies where comprehensive screening of a wide range of phytohormones is desired [4]. We describe in this article our method of choice for comprehensive phytohormone extraction and HPLC-separation including all the pitfalls and difficulties that may occur during method establishment and how to address these.

Most of the publications rely on the multi reaction monitoring (MRM) mass spectrometric detection, using compound specific precursor-to-transitions performed on triple quad mass spectrometers [5,6,7]. This approach is popular due to high sensitivity and selectivity which is achieved by reduced matrix background. In contrast, relatively few studies describe the use of high-resolution mass spectrometry (HRMS) for phytohormone analysis. We compared both approaches and describe the advantages and limitations of both detection strategies for comprehensive phytohormone analysis.

## Method details

### Phytohormone Classes

In this publication we show the parallel analysis of the phytohormone classes: auxins, cytokinins, gibberellins, ABA, SA and jasmonates. The major bioactive forms of auxins and cytokinins are indole-acetic acid (IAA) and *trans*-zeatin. Sometimes the terms “auxin” and “cytokinin” are used as synonyms for these phytohormones. We strongly recommend describing exactly which phytohormone molecules are analyzed in a study to enable readers to correctly interprete the data and compare data between publications. For example, the term “zeatin” can refer to the bioactive *trans*-zeatin, but also to its isomer *cis*-zeatin. In addition, both of these cytokinins are present in far higher concentrations in various glycosylated forms. Furthermore, the structurally related cytokinins dihydrozeatin and isopentenyladenine (IP) exist. Thus, the information which cytokinin molecules were actually quantified is crucial to understand the data. This holds also true for the other phytohormone classes.

IAA can be present as a methyl- or ethyl-ester or it can be conjugated to a number of different amino acids. Gibberellins (GAs) are tetracyclic diterpene acids with over 100 different members of this group discovered so far. Only a few of the many described GAs have so far been shown to be bioactive, namely GA1, GA3, GA4 and GA7. GA4 is the major bioactive form in the model plant Arabidopsis, but others may dominate in other plant species. ABA can be present as different isomers and also in a conjugated glycosylated form. The phytohormone class jasmonates is synthesized from linolenic acid and comprises jasmonic acid (JA) isomers and their methyl esters as well as their biosynthetic precursor *cis*-12-oxo-phytodienoic acid (*cis*-OPDA) and amino acid conjugates of JA such as JA-isoleucine. In addition, JA metabolites include various modified forms of JA, e.g. as a result of hydroxylation or glucosylation, such as 12-OH-JA and 12-O-Glc-JA.

We compared existing methods for the parallel analysis of six phytohormone classes (auxins, cytokinins, GAs, ABA, SA, and jasmonates, Figure 1) and developed optimization steps with regard to extraction, HPLC separation and mass spectrometry analysis. In order to help other groups, and especially newcomers to the field, we would like to share our findings, including optimizations and possible pitfalls that might occur.

**Figure 1:**
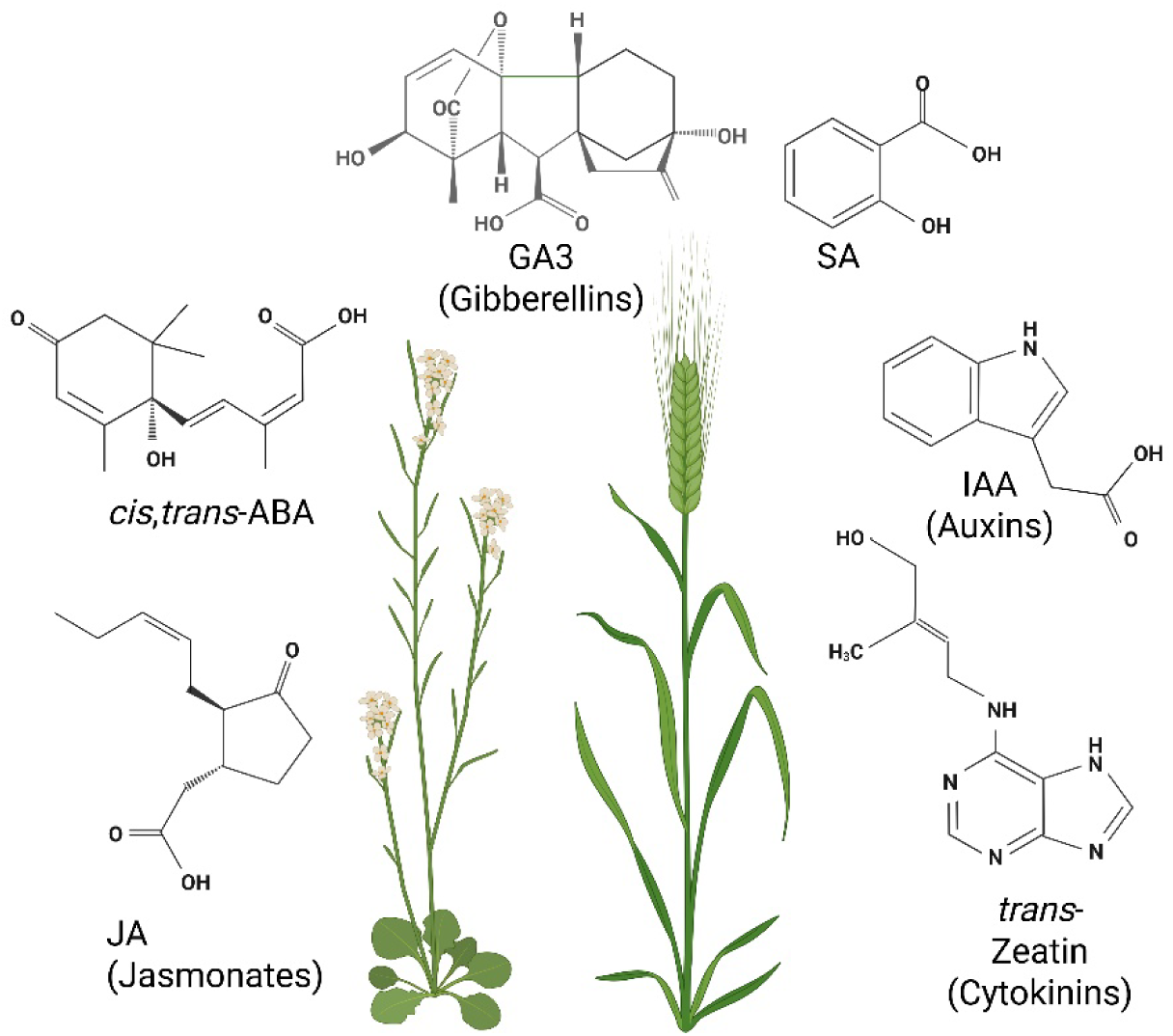
Phytohormone classes analyzed in this study: jasmonates, abscisic acid (ABA), gibberellins, salicylic acid (SA), auxins and cytokinins. Depicted are representative phytohormone structures and the model plant species *Arabidopsis thaliana* as well as the crop plant barley (*Hordeum vulgare*).

Parallel quantification of different phytohormone classes is very challenging due to the strongly different physicochemical properties between the different classes of phytohormones, but also within a phytohormone class. This affects solubility in different solvents, compound stability and interactions with other compounds.

The concentrations of phytohormones can range from extremely low concentrations to strong accumulations of some phytohormones under certain conditions. Phytohormone levels are dependent e.g. on plant species, organ, tissue and developmental stage and on external stress factors such as temperature, drought or biotic stress.

We are often confronted with biological questions where the amount of plant material that is available for phytohormone analysis is limited. Thus, it is not always possible to increase the amount of plant material to enrich and purify phytohormones which are present at low concentrations. Limitations in plant material also prevent us from choosing for each phytohormone class different extraction methods, which would mean several separate extractions thus multiplying sample numbers.

This dilemma of limited availability of plant material requires a simple but comprehensive extraction method in which as many different phytohormones as possible are extracted. In a second step a robust and powerful HPLC separation of the extracted phytohormones is required to allow confident identification of phytohormones and separation of isomers. Finally, the MS detection method has to be carefully chosen to meet the demands of the study. To this end we compared the advantages and disadvantages of HRMS versus MRM and describe the optimization steps we implemented to ensure high sensitivity and selectivity.

The optimal method should provide a broad overview of the phytohormone status of a given plant tissue, knowing that a parallel analysis of the different phytohormone classes in a complex plant extract is always a compromise. The method we describe will be well-suited to monitor changes in phytohormones between tissues, growth conditions or mutants.

When choosing the detection strategy, it is important to tailor it to the biological question at hand. A different strategy can be suitable when assessing the phytohormone status of a non-model plant in contrast to targeting a specific set of well-known phytohormones in a model plant such as *Arabidopsis thaliana*. In non-model plants little may be known about the specific phytohormones within a given phytohormone class. Among the gibberellins, for example, there is a high chemical diversity in possible structures, therefore the analysis of gibberellins in a non-model plant can resemble a semi-untargeted approach.

With regard to detection, we used two strategies:

- High-resolution mass spectrometry (HRMS), which allows for the determination of an accurate mass up to several decimal places and thus enables the identification of phytohormone ions in a matrix without fragmentation.

In this study we employed a Q-TOF instrument with a resolution of 60.000, which can be exceeded by modern multi-reflecting TOF instruments or Orbitrap instruments reaching up to over 100.000. HRMS is usually the method of choice for untargeted screening of metabolites.

- Tandem mass spectrometry employing compound-specific precursor-to-product transitions (multiple reaction monitoring = MRM) on a triple quadrupole instrument.

This method relies on the selection of a phytohormone “precursor ion” with a certain mass (or mass-to-charge ratio, m/z) in the first quadrupole. Subsequently the selected precursor ions are fragmented by collision with an inert gas. The third quadrupole is set to a certain mass window selectively letting only specific “product ions” pass through, which are then detected. The combination of the mass of precursor ion and the mass of the product ion is called “precursor-to-product transition” or “precursor-product ion pair” and sometimes shortly referred to as “MRM” or “transition”. This method is known to be highly specific and sensitive and is most-widely used in plant hormone analysis. Here we used a QTrap instrument.

### Reagents and Chemicals

Standards were purchased from OlChemim Ltd. (Czech Republic) except for SA and IAA which were purchased from Sigma-Aldrich (Germany) and Supelco (Germany). Isotopically labeled standards were purchased whenever possible but these were not always commercially available or out of stock. Methanol for LC-MS analysis was purchased from Biosolve (The Netherlands) in ULC/MS grade.

### Internal Standards

For a most reliable absolute quantification of phytohormones, the addition of suitable isotope-labeled internal standards (I.S.) is extremely helpful. Isotope-labeled standards are added to a phytohormone extract at a defined concentration. Due to the fact that these standards are chemically almost identical to the corresponding phytohormones, they also behave almost identically during extraction, HPLC separation and ionization. At the same time they can be distinguished during MS analysis due to their deviating mass. We recommend using as many suitable isotope-labeled standards as possible, to cover every phytohormone species. However, if suitable isotope-labeled standards are not available for all investigated phytohormones, related isotope-labeled standards or other related phytohormone standards from the same phytohormone class which do not naturally occur in the investigated plant tissues can be used as an alternative. In this study we used e.g. the isotope-labeled standard ^2^H_2_-GA9 to quantify GA9 as well as GA3 because no isotope-labeled standard was available for GA3. An example for a not isotopically labeled standard is 9,10-dihydrojasmonic acid, which has been used as a standard for the analysis of jasmonates.

To obtain meaningful results it is important to choose an optimal concentration of internal standards. We empirically tested this and arrived at 75 nM final concentration in a final volume of 100 µl sample extractor the following reasons: Most of the available deuterated standards have a minimum 95 % purity, meaning that a minor contamination of the standard with unlabeled phytohormones cannot be excluded. We recommend testing every labeled standard before use to assess the degree of contaminations. Due to the very low levels of naturally occurring phytohormones it is important to ensure that possible traces of unlabeled phytohormones originating from the internal standard mix remain below the limit of detection in the final sample extracts.

Furthermore, the internal standards work best as a reference for the naturally occurring phytohormones when they are used in a physiological concentration range. For some very expensive and precious isotope-labeled standards, such as ^2^H_6_-JA, we used a lower concentration of 25 nM in some samples.

### Harvest of Plant Material

To avoid phytohormone degradation we recommend using only fresh plant tissue, harvested directly prior to extraction and stored in liquid nitrogen, on dry ice or at -80°C for as little time as possible. Depending on which phytohormones should be analyzed, different aspects have to be considered when the harvest of tissue samples is planned.

There are publications on the circadian fluctuation of some phytohormones [8]. This shows that the timepoint of harvest during the day can severely influence the phytohormone level. When large numbers of samples are harvested, a randomized harvesting is therefore strongly recommended to protect against an artificial bias introduced by daytime.

Other phytohormones such as jasmonates can accumulate very quickly upon wounding, therefore a very careful handling of the plants is strongly recommended. This includes rapid freezing of the harvested plant tissues in liquid nitrogen to immediately stop any enzymatic processes in the cells.

### Phytohormone Extraction from Plant Material - Liquid-Liquid Extraction versus Solid Phase Extraction

For the parallel extraction of different phytohormone classes from plants we have tested different existing protocols. We compared a widely-used simple extraction method which uses just one single cleanup step by liquid-liquid extraction (LLE) to a more elaborate extraction using a solid phase extraction (SPE) column to separate acidic and alkaline phytohormone classes (Figure 2). The methods we compared were based on protocols first described by Pan et al. [5,9] and Eggert and von Wiren [10], respectively. The liquid-liquid extraction protocol uses a mixture of isopropanol and water, acidified with HCl to extract phytohormones from homogenized plant tissues. In a second step phytohormones are separated from other compounds by phase separation against dichloromethane (DCM). In this step most phytohormones locate primarily to the DCM phase, which is collected and concentrated under a stream of nitrogen. No further clean-up or enrichment steps are implemented.

The other extraction protocol we assessed is loosely based on the clean-up of phytohormones described in [10]. We adapted parts of this method to include one comprehensive extraction for all phytohormone classes using 80% MeOH followed by re-extraction with 50% MeOH. The combined extracts were applied to SPE on an MCX column (Waters). Phytohormones were eluted from the column with 90% acetonitrile and 0.1% formic acid (FA) (for acidic phytohormones) and subsequently with 60% acetonitrile containing 5% ammonia (for alkaline phytohormones).

**Figure 2:**
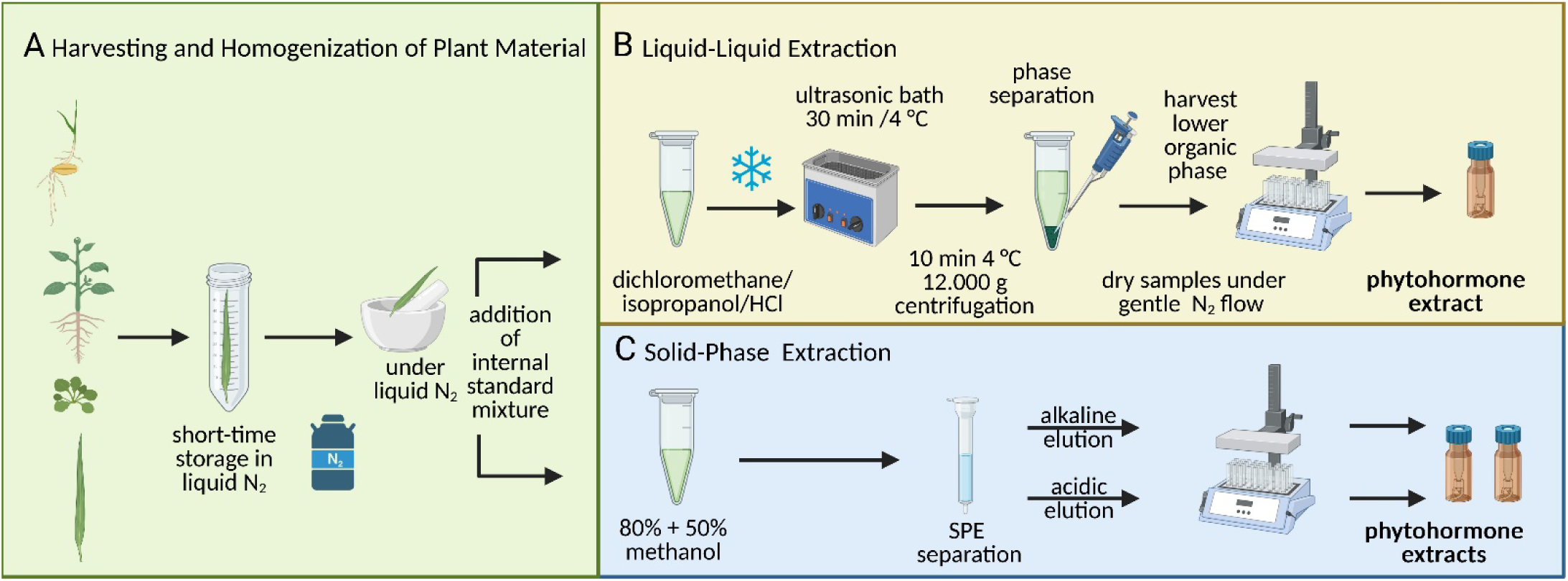
Workflow for comparison of phytohormone extraction protocols. **A**: Harvesting and homogenization of plant material. **B**: Phytohormone extraction using liquid-liquid extraction. **C**: Phytohormone extraction with 80% and 50% methanol and subsequent purification by solid phase extraction (SPE). SPE is used to separate alkaline and acidic phytohormones yielding two separate phytohormone fractions for further analysis.

We evaluated the recovery of a mixture of reference standards, including a variety of cytokinins, IAA, ABA, and gibberellins, using LLE method versus the SPE clean-up method.

Among the alkaline phytohormones free zeatin (*cis*- and *trans*-zeatin) as well as dihydrozeatin showed a higher recovery rate using LLE than SPE. The more polar conjugated cytokinins, *cis*-zeatin-riboside, *trans*-zeatin-riboside and especially *trans*-zeatin-9-glucoside showed a higher recovery rate using SPE. In contrast, IP showed a much better recovery using LLE, while isopentenyladenosine (IPR) worked equally well using either method. The acidic phytohormones, gibberellins, ABA as well as IAA, showed a very good recovery using either method, with a minor advantage for gibberellins when LLE was used.

Taken together the only phytohormones for which we observed an advantage in the more elaborate SPE-based extraction were the glucosylated zeatins. Most likely due to their relatively high polarity we saw significant losses of these during LLE, which was not observed in the other method. However, using isotope-labeled standards their quantification is still possible because glucosylated zeatins are often present at much higher concentrations than the bioactive free form *trans*-zeatin.

The method including the SPE clean-up is far more time-consuming than the LLE. Correct adjustment of pH and application of large volumes of elution solutions, which later have to be evaporated, make the handling of the SPE method more complicated and harder to transfer between laboratories.

We would further like to address the general problem of additional purification steps and the use of SPE columns, which present the danger of phytohormones sticking to the surfaces, leading to loss in the process. Furthermore, all SPEs have to be carefully chosen and pre-cleaned to minimize the introduction of plasticizers or other contaminants into the mass spectrometer.

It has to be mentioned that we often deal with limited amounts of plant tissue, e.g. when working with plant meristems. This, combined with the extremely low concentrations of some phytohormones makes it even more important to minimize losses due to surface adhesion. Limited amounts of plant material also contradict the major useful function of SPE, namely the enrichment of phytohormones from large amounts of plant tissue.

Taken together we found LLE to be best suited for our requirements: a fast and comprehensive extraction method which yields a broad range of gibberellins, cytokinins, auxins, ABA, SA and jasmonates. The method is very suitable and we modified the method to optimize the procedure for our needs. When Pan et al. first described their method they used it to target 18 different phytohormones in the model plant Arabidopsis, focusing on the major Arabidopsis phytohormones and their methylated forms [5, 9]. We have optimized the method for the analysis of about 50 different phytohormones using standards, focusing also on conjugated forms of cytokinins and IAA. We applied our optimized method for the analysis of different tissues from the model plant Arabidopsis, as well as crop plants like rice, barley and tobacco [11, 12].

We would like to highlight the following modified steps for optimization:

- To obtain the finest possible powder we recommend grinding approx. 50 mg of plant material in a porcelain mortar under liquid nitrogen and then weighing the material quickly on a sensitive balance without thawing.
- To minimize loss of plant material in cases where as little as 5-10 mg plant material was available we harvested it in 1.5 mL reaction tubes and homogenized the material to a fine powder under liquid nitrogen using a polypropylene pistil inside the reaction tube. We then added some extraction solution and continued grinding, finally rinsing the pistil with some fresh extraction solution.
- To minimize phytohormone loss due to surface adhesion we used Eppendorf® Protein-LoBind reaction tubes throughout the process. We found these high-quality reaction tubes superior to other tubes both with regard to low metabolite adhesion and low plasticizer leakage.
- To ensure that the isotope-labeled standards are present throughout the extraction process we added the extraction solution including the internal standard mix directly after grinding. In some publications the internal standards are added at a later step [10], which we do not recommend. The function of internal standards is to reflect any changes in signal intensity of phytohormones occurring during the extraction process and during subsequent storage and analysis. Among others this can include degradation of phytohormones over time, effects of temperature and pH, solubility in different solvents etc. Furthermore, any variation in sample treatment such as evaporation of solvents or pipetting errors will be reflected by the internal standard. Therefore, the internal standards should be subjected to the same treatment as the phytohormones in the plant extract.
- As a control for the extraction efficiency and the integrity of the internal standard mixture we always prepared quality controls containing only the internal standard mix without plant material. These controls are submitted to the exact same extraction procedure and are measured in parallel to the plant samples.
- To further improve extraction efficiency, we used an ice-cooled ultrasonic bath instead of a shaker during the two extraction steps. This was done once after addition of the extraction solution and then again after addition of DCM. During these 30 minutes we also vortexed the samples several times. In our experience ultrasonic baths increase extraction efficiencies and are a good alternative to extractions on a shaker overnight which are sometimes recommended. A quick extraction also minimizes the risk of phytohormone degradation over time. The ultrasonic bath can be filled partly with water and partly with ice floating around to ensure that the temperature does not exceed 4 °C.
- At the end of the extraction process the samples are dried under a stream of N2. In [5] it is mentioned that the samples should be concentrated to almost dryness and then re-dissolved in 100% MeOH. We followed this protocol and realized that it can be difficult to ensure that all samples are removed from the sample concentrator just in time before they reach dryness, because not all samples require exactly the same time to dry. We checked all samples regularly and removed those samples from the concentrator that had just reached dryness. We took care to avoid prolonged exposition to the stream of N_2_ after dryness because this makes it more difficult to re-dissolve phytohormones.
- To improve HPLC separation of phytohormone extracts, we changed the final solvent to re-dissolve the dried samples. The dried extracts are readily soluble in 100% MeOH. However, this solvent has a high eluting power. When used as a sample solvent this can be disadvantageous for a good HPLC separation. In our hands the injection of 10 µl sample in 100% MeOH resulted in premature elution of many cytokinins from the HPLC column, leading to split peaks and loss in signal intensity. This could be overcome by adding H_2_O to the samples to a final ratio of MeOH/H_2_O 1:1. Hence, we dissolved all dried samples first in 50 µl 100% MeOH and then added another 50 µl of H_2_O.
- While the dried extracts are readily soluble in 100% MeOH, the addition of H_2_O resulted in some insoluble precipitates, which had to be removed prior to LC-MS analysis. Precipitation was mostly observed after keeping the samples in a cooled autosampler for some time. To protect our HPLC column and prevent blockage we added an additional step where we kept the samples at 4 °C overnight followed by a 10-minute centrifugation step at high speed. We collected the supernatant which was then used for LC-MS analysis. Analysis of our internal standards showed that no losses in signal intensities were observed by this procedure.
- Instead of sample filtration prior to the LC separation to protect the HPLC columns, we prefer to use vendor-recommended guard columns. In case of contamination or blockage these can be exchanged at a lower price than exchanging the HPLC column. Thus, we can avoid the introduction of an additional step where contaminants may be introduced or phytohormones may be lost due to surface adhesion.

### Chromatographic Separation of Phytohormones

In addition to the extraction method, we optimized the HPLC method for chromatographic separation of a large number of phytohormones. We would like to highlight the importance of an optimized HPLC method for separating phytohormone isomers.

Phytohormones were separated on a Dionex 3000 HPLC system (Thermo Scientific, Germany) with a binary gradient system and analyzed on a high-resolution Q-TOF mass spectrometer (maXis 4G, Bruker Daltonics, Germany).

We tested several HPLC columns for optimal separation of all phytohormone classes. A C18 XSelect™ HSST3 Column, 2.5 µm, 3 mm X 150 mm (Waters, Ireland) and a C18 Atlantis™ Premier BEH column, 2.5 µm, 2.1 mm X 150 mm (Waters, Ireland) were selected as the best candidates. The HSST3 Column is recommended by the manufacturer as an ideal column for the parallel separation of polar and nonpolar metabolites. As phytohormones comprise polar metabolites such as cytokinins as well as nonpolar metabolites such as jasmonates this was an important criterion to choose this column. The Atlantis column is recommended especially for a reverse-phase separation of polar acidic metabolites.

Both HPLC columns are compatible with 100% organic solvent which was a benefit for our gradient but also for column maintenance. Phytohormone extracts contain additional nonpolar substances which are co-extracted from plant material along with the phytohormones and which might contaminate the HPLC column over time. Compatibility with 100 % organic solvent is advantageous for regular column clean-up routines to remove such substances from the column.

On both columns phytohormones were separated using a binary gradient system with mobile phase A 100% H_2_O + 0.1% FA and mobile phase B 100% methanol + 0.1% FA. To ensure a prolonged separation of polar phytohormones we started the gradient at 2% mobile phase B, and increased it slowly to 99% B from 1-18 min. To ensure enough time for nonpolar metabolites to elute from the column the plateau was held at 99% for 4 min. The gradient was returned to 2% mobile phase B within 1 min. Prior to the next injection the system was equilibrated at 2% B for 7 min. To optimize separation as well as solvent use and backpressure, we chose a flow rate of 0.35 ml/min.

Tables 1 and 2 show the full set of standards analyzed in this study by high-resolution Q-TOF MS, including the [M+H]^+^/ [M-H]^-^ in positive or negative ionization mode and their retention times on the Waters HSST3 (Table 1) or Waters Atlantis (Table 2) HPLC column with the described LC gradient.

**Table 1:** Retention times [min] and accurate masses [M-H]^-^ / [M+H]^+^ of 52 phytohormone standards eluting from the Waters HSST3 HPLC column and measured by Q-TOF MS in their respective optimal ionization modes (32 neg/20 pos).

| Standard | [M-H]<sup>-</sup> (negative mode) | RT [min] |
| --- | --- | --- |
| ABA |  |  |
| (±)-*cis,trans*-Absciscic acid (ct-ABA) | [M-H]<sup>-</sup> = 263.129 | 14.6 |
| I.S. <sup>2</sup>H<sub>6</sub> (+)-*cis,trans*-Absciscic acid (<sup>2</sup>H<sub>6</sub>-ABA) | [M-H]<sup>-</sup> = 269.167 | 14.5 |
| (+)-*trans,trans*-Absciscic acid (S-tt-ABA) | [M-H]<sup>-</sup> = 263.129 | 14.1 |
| Gibberellins |  |  |
| Gibberellin A1 (GA1) | [M-H]<sup>-</sup> = 347.150 | 12.9 |
| I.S. <sup>2</sup>H<sub>4</sub>-Gibberellin A1 (<sup>2</sup>H<sub>4</sub>-GA1) | [M-H]<sup>-</sup> = 351.175 | 12.7 |
| Gibberellic Acid (GA; GA3) | [M-H]<sup>-</sup> = 345.134 | 12.5 |
| Gibberellin A4 (GA4) | [M-H]<sup>-</sup> = 331.155 | 17.1 |
| I.S. <sup>2</sup>H<sub>2</sub>-Gibberellin A4 (<sup>2</sup>H<sub>2</sub>-GA4) | [M-H]<sup>-</sup> = 333.168 | 17.1 |
| Gibberellin A8 (GA8) | [M-H]<sup>-</sup> = 363.145 | 10.6 |
| Gibberellin A9 (GA9) | [M-H]<sup>-</sup> = 315.160 | 18.0 |
| I.S. <sup>2</sup>H<sub>2</sub>-Gibberellin A9 (<sup>2</sup>H<sub>2</sub>-GA9) | [M-H]<sup>-</sup> = 317.173 | 18.0 |
| Gibberellin A19 (GA19) | [M-H]<sup>-</sup> = 361.166 | 16.1 |
| I.S. <sup>2</sup>H<sub>2</sub>-Gibberellin A19 (<sup>2</sup>H<sub>2</sub>-GA19) | [M-H]<sup>-</sup> = 363.178 | 16.0 |
| Gibberellin A20 (GA20) | [M-H]<sup>-</sup> = 331.155 | 15.3 |
| I.S. <sup>2</sup>H<sub>2</sub>-Gibberellin A20 (<sup>2</sup>H<sub>2</sub>-GA20) | [M-H]<sup>-</sup> = 333.168 | 15.3 |
| Gibberellin A34 (GA34) | [M-H]<sup>-</sup> = 347.150 | 16.2 |
| I.S. <sup>2</sup>H<sub>2</sub>-Gibberellin A34 (<sup>2</sup>H<sub>2</sub>-GA34) | [M-H]<sup>-</sup> = 349.163 | 16.1 |
| Gibberellin A44 (GA44) | [M-H]<sup>-</sup> = 345.171 | 15.8 |
| I.S. <sup>2</sup>H<sub>2</sub>-Gibberellin A44 (<sup>2</sup>H<sub>2</sub>-GA44) | [M-H]<sup>-</sup> = 347.183 | 15.8 |
| Gibberellin A53 (GA53) | [M-H]<sup>-</sup> = 347.186 | 17.3 |
| I.S. <sup>2</sup>H<sub>2</sub>-Gibberellin A53 (<sup>2</sup>H<sub>2</sub>-GA53) | [M-H]<sup>-</sup> = 349.199 | 17.3 |
| Salicylic Acid and Others |  |  |
| Salicylic acid (SA) | [M-H]<sup>-</sup> = 137.024 | 14.4 |
| I.S. <sup>2</sup>H<sub>4</sub>-Salicylic acid (<sup>2</sup>H<sub>4</sub>-SA) | [M-H]<sup>-</sup> = 141.050 | 14.3 |
| Cinnamic acid (CA) | [M-H]<sup>-</sup> = 147.045 | 15.4 |
| Jasmonates |  |  |
| (±)-Jasmonic acid (JA) | [M-H]<sup>-</sup> = 209.118 | 15.8 |
| I.S. <sup>2</sup>H<sub>6</sub>-(±)-Jasmonic acid (<sup>2</sup>H<sub>6</sub>-JA) | [M-H]<sup>-</sup> = 215.156 | 15.8 |
| I.S. (±)-9,10-Dihydrojasmonic acid (9,10-DHJA) | [M-H]<sup>-</sup> = 211.134 | 16.5 |
| N-[(-)-Jasmonoyl]-(S)-isoleucine (Jaile) | [M-H]<sup>-</sup> = 322.202 | 17.2 |
| N-[(-)-Jasmonoyl]-(L)-isoleucine (Jaile) | [M-H]<sup>-</sup> = 322.202 | 17.2 |
| I.S. <sup>2</sup>H<sub>6</sub>-N-[(-)-Jasmonoyl]-(L)-isoleucine (<sup>2</sup>H<sub>6</sub>-Jaile) | [M-H]<sup>-</sup> = 328.240 | 17.1 |
| *cis*-12-Oxo-Phytodienoic acid (*cis*-OPDA) | [M-H]<sup>-</sup> = 291.197 | 18.9 |
| I.S. <sup>2</sup>H<sub>5</sub>-*cis*-12-Oxo-Phytodienoic acid (<sup>2</sup>H<sub>5</sub>-OPDA) | [M-H]<sup>-</sup> = 296.228 | 18.9 |
| Standard | [M+H]<sup>+</sup> (positive mode) | RT [min] |
| Jasmonates |  |  |
| (±)-Jasmonic acid methyl ester (MeJA) | [M+H]<sup>+</sup> = 225.149 | 17.3 |
| Auxins |  |  |
| Indole-3-acetic acid (IAA) | [M+H]<sup>+</sup> = 176.071 | 13.6 |
| I.S. <sup>2</sup>H<sub>5</sub>-Indole-3-acetic acid (<sup>2</sup>H<sub>5</sub>-IAA) | [M+H]<sup>+</sup> = 181.102 | 13.5 |
| Indole-3-acetyl-aspartic acid (IAAasp) | [M+H]<sup>+</sup> = 291.098 | 11.7 |
| Indole-3-acetyl-glutamic acid (IAGlu) | [M+H]<sup>+</sup> = 305.113 | 11.9 |
| Indole-3-acetyl-alanine (IAAla) | [M+H]<sup>+</sup> = 247.108 | 12.8 |
| Indole-3-acetyl-valine (IAVal) | [M+H]<sup>+</sup> = 275.139 | 14.9 |
| Indole-3-acetic acid methyl ester (IAAME) | [M+H]<sup>+</sup> = 190.086 | 15.2 |
| Indole-3-acetyl-tryptophane (IATrp) | [M+H]<sup>+</sup> = 362.150 | 15.6 |
| Indole-3-acetyl-isoleucine (IAIleu) | [M+H]<sup>+</sup> = 289.155 | 15.7 |
| Indole-3-acetyl-phenylalanine (IAPhe) | [M+H]<sup>+</sup> = 323.139 | 16.0 |
| Indole-3-acetic acid ethyl ester (IAAEt) | [M+H]<sup>+</sup> = 204.102 | 16.3 |
| Indole-3-acetamide (IAM) | [M+H]<sup>+</sup> = 175.087 | 11.5 |
| Indole-3-propionamide (IPAM) | [M+H]<sup>+</sup> = 189.102 | 12.7 |
| Indole-3-carboxylic acid (I3CA) | [M+H]<sup>+</sup> = 162.055 | 13.2 |
| Indole-3-carboxylic acid methyl ester (I3CAME) | [M+H]<sup>+</sup> = 176.071 | 15.4 |
| Indole-3-acetonitrile (I3ACN) | [M+H]<sup>+</sup> = 157.076 | 14.0 |
| I.S. <sup>2</sup>H<sub>4</sub>-Indole-3-acetonitrile (<sup>2</sup>H<sub>4</sub>-I3ACN) | [M+H]<sup>+</sup> = 161.101 | 13.9 |
| Indole-3-propionic acid (IPA) | [M+H]<sup>+</sup> = 190.086 | 14.6 |
| Indole-3-butyric acid (IBA) | [M+H]<sup>+</sup> = 204.102 | 15.4 |

I.S. = internal standard; <sup>2</sup>H<sub>x</sub>= deuterated with x deuterium atoms

**Table 2:** Retention times [min] and accurate masses [M+H]^+^ of 20 cytokinin standards eluting from the Waters Atlantis HPLC column and measured by Q-TOF MS in positive ionization mode.

| Standard | [M+H]<sup>+</sup> (positive mode) | RT [min] |
| --- | --- | --- |
| Cytokinins |  |  |
| *trans*-Zeatin (tZ) | [M+H]<sup>+</sup> = 220.119 | 6.0 |
| I.S. <sup>2</sup>H<sub>5</sub>-*trans*-Zeatin (<sup>2</sup>H<sub>5</sub>-tZ) | [M+H]<sup>+</sup> = 225.151 | 6.0 |
| *cis*-Zeatin (cZ) | [M+H]<sup>+</sup> = 220.119 | 6.4 |
| I.S. <sup>15</sup>N<sub>4</sub>-*cis*-Zeatin (<sup>15</sup>N<sub>4</sub>-cZ) | [M+H]<sup>+</sup> = 224.107 | 6.4 |
| Dihydrozeatin (DHZ) | [M+H]<sup>+</sup> = 222.135 | 6.0 |
| I.S. <sup>2</sup>H<sub>5</sub>-*trans*-Zeatin-O-glucoside (<sup>2</sup>H<sub>5</sub>-tZOG) | [M+H]<sup>+</sup> = 387.203 | 5.9 |
| I.S. <sup>2</sup>H<sub>5</sub>-*trans*-Zeatin-7-glucoside (<sup>2</sup>H<sub>5</sub>-tZ7G) | [M+H]<sup>+</sup> = 387.203 | 5.9 |
| *cis*-Zeatin-O-glucoside (cZOG) | [M+H]<sup>+</sup> = 382.172 | 6.2 |
| *trans*-Zeatin-9-glucoside (tZ9G) | [M+H]<sup>+</sup> = 382.172 | 6.9 |
| I.S. <sup>2</sup>H<sub>5</sub>-*trans*-Zeatin-9-glucoside (<sup>2</sup>H<sub>5</sub>-tZ9G) | [M+H]<sup>+</sup> = 387.203 | 6.8 |
| *cis*-Zeatin-9-glucoside (cZ9G) | [M+H]<sup>+</sup> = 382.172 | 7.2 |
| I.S. <sup>15</sup>N<sub>4</sub>-*cis*-Zeatin-9-glucoside (<sup>15</sup>N<sub>4</sub>-cZ9G) | [M+H]<sup>+</sup> = 386.160 | 7.2 |
| I.S. <sup>2</sup>H<sub>5</sub>-*trans*-Zeatin-O-glucoside riboside (<sup>2</sup>H<sub>5</sub>-tZROG) | [M+H]<sup>+</sup> = 519.246 | 8.1 |
| *cis*-Zeatin-O-glucoside riboside (cZROG) | [M+H]<sup>+</sup> = 514.214 | 8.5 |
| *trans*-Zeatin riboside (tZR) | [M+H]<sup>+</sup> = 352.162 | 8.7 |
| *cis*-Zeatin riboside (cZR) | [M+H]<sup>+</sup> = 352.162 | 9.1 |
| N6-Isopentenyladenine (IP) | [M+H]<sup>+</sup> = 204.124 | 9.3 |
| I.S. <sup>15</sup>N<sub>4</sub>-N6-Isopentenyladenine (<sup>15</sup>N<sub>4</sub>-IP) | [M+H]<sup>+</sup> = 208.113 | 9.3 |
| N6-Isopentenyladenosine (IPR) | [M+H]<sup>+</sup> = 336.167 | 11.9 |
| I.S. <sup>2</sup>H<sub>6</sub>-N6-Isopentenyladenosine (<sup>2</sup>H<sub>6</sub>-IPR) | [M+H]<sup>+</sup> = 342.204 | 11.9 |

I.S. = internal standard; <sup>2</sup>H<sub>x</sub>= deuterated with x deuterium atoms

Our final recommendation for the parallel LC-MS analysis of different phytohormone classes is to use the Waters HSST3 column. Figure 3 shows the separation of phytohormone standards (auxins, gibberellins, SA, ABA and jasmonates) on the Waters HSST3 column. Phytohormones were measured by Q-TOF MS in both positive and negative ionization mode separately. For most phytohormones, an optimal ionization mode was determined and subsequently used for further analyses. The optimal ionization mode for each phytohormone can be found in Table 1. Figure 3 shows the extracted ion chromatograms (EICs) for 48 standards of the phytohormone classes auxins (depicted in brown), gibberellins (green), SA (purple), ABA (yellow) and jasmonates (blue) measured under their respective optimal ionization conditions. EICs of jasmonic acid-isoleucine (JA-Ileu, both S- and L-isomers) and OPDA are depicted in Figure 3A, measured in negative ionization mode, and in Figure 3B, measured in positive ionization mode, because they can be detected equally well under both ionization conditions.

**Figure 3:**
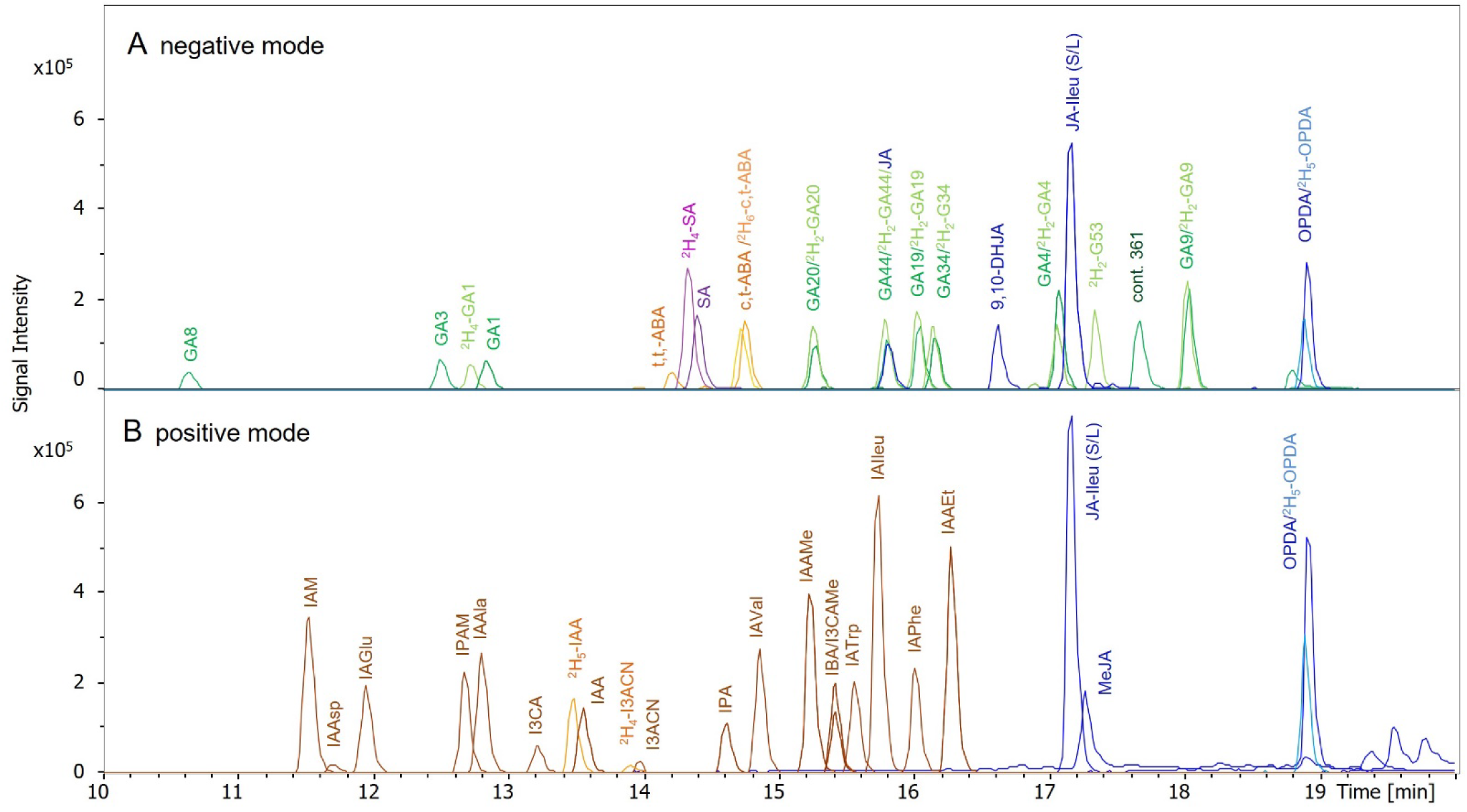
Chromatographic separation of a mixture of 48 phytohormone standards on the Waters HSST3 HPLC column. Depicted are the extracted ion chromatograms (EICs) of the respective [M-H]^-^ / [M+H]^+^ in their optimal ionization modes. **A**: Analysis in negative ionization mode. Gibberellins are depicted in green, SA in purple, ABA in yellow and jasmonates in blue. Cont. 361 refers to a consistently observed contaminant with an m/z of 361.166, which is isobaric to GA19 and therefore shows up on the EIC. **B**: Analysis in positive ionization mode. IAA and other auxins are depicted in brown and jasmonates in blue. The respective isotope-labeled standards are depicted in lighter shades of the same colors. Some phytohormones, which are equally well ionized in negative or positive ionization mode are shown in both A and B. All standards were dissolved at a concentration of 75 nM in MeOH/H_2_O (except ^2^H_4_-SA at 150 nM).

For special applications such as the distinct quantification of different zeatin-glucoside isomers we recommend using the Waters Atlantis column (Figure 5).

For many phytohormones stereoisomers have been observed in plant tissues, e.g. *cis*/*tran*s-isomers are very common. These stereoisomers may differ profoundly in their biological activity. Thus, in order to correctly interpret the biological impact of phytohormone level changes in a given experiment, it is important to distinguish between these isomers during phytohormone analysis. These isomers are not distinguishable by accurate mass alone and can also display an almost identical fragmentation behavior. Therefore, it is very important to optimize the chromatographic separation of phytohormone extracts with regard to the separation of the frequently occurring isomers.

Prominent examples for phytohormone stereoisomers are *cis*-zeatin and *trans*-zeatin, which are known to differ in their biological function and are well-separated on both the Atlantis and the HSST3 HPLC column. *Trans*- and *cis*-zeatin form the basis for a variety of conjugated cytokinins, such as *trans*- and *cis*-zeatin-riboside. For *trans*- and *cis*-zeatin-glucoside, the glucose can be attached as an N-glucoside in position 7 or 9 or as an O-glucoside, giving rise to six different zeatin-glucoside isomers: *cis*-zeatin-O-glucoside (cZOG), *trans*-zeatin-O-glucoside (tZOG), *cis*-zeatin-9-glucoside (cZ9G), *trans*-zeatin-9-glucoside (tZ9G), *cis*-zeatin-7-glucoside (cZ7G) and *trans*-zeatin-7-glucoside (tZ7G) (Figure 4).

**Figure 4:**
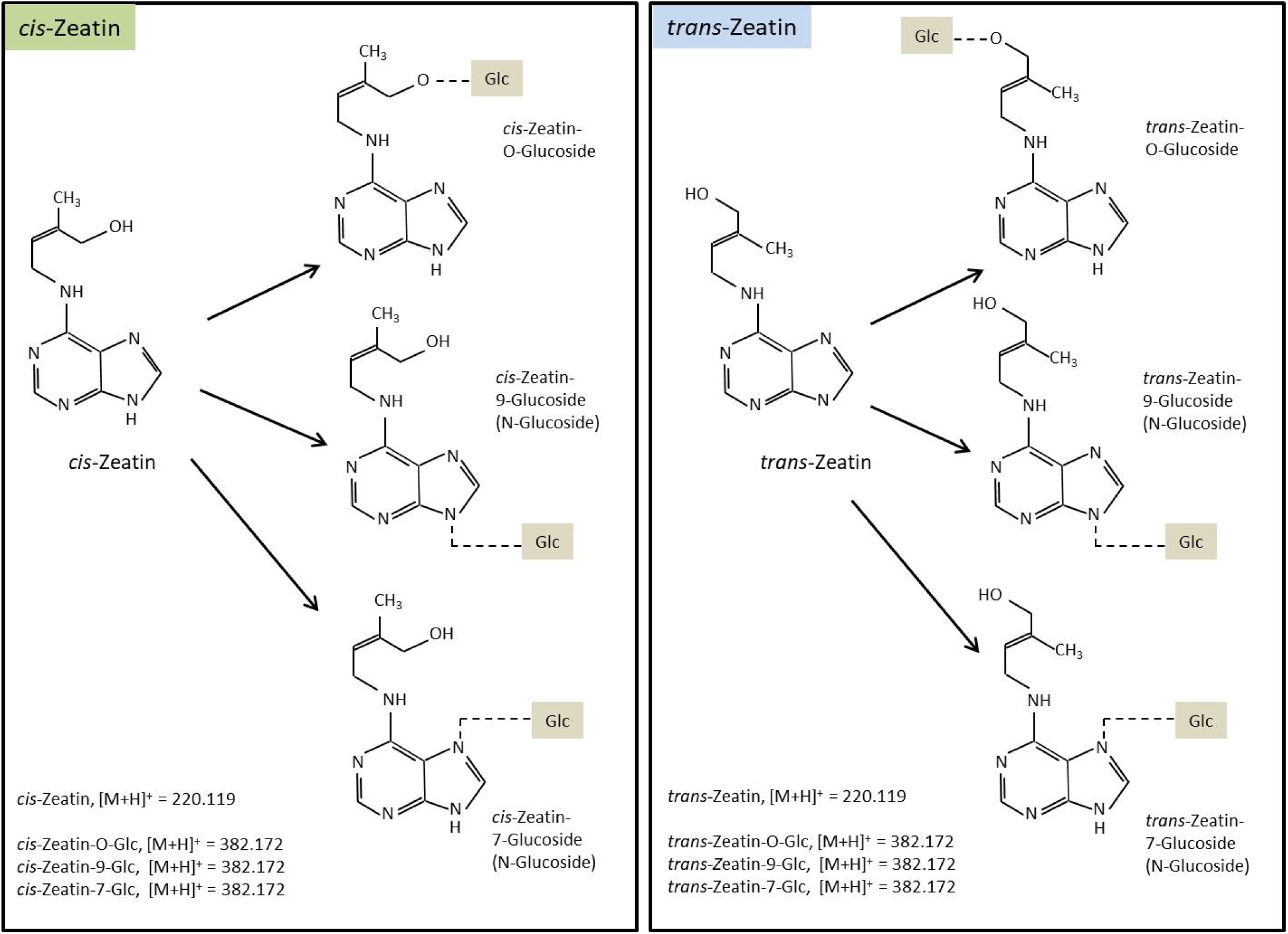
Free zeatin and zeatin-glucoside isomers in plants. Free zeatin can occur as *cis*-zeatin or *trans*-zeatin. N-glucosylation can take place at one of the nitrogen atoms at position 7 or 9 of the zeatin ring system. O-glucosylation occurs at the free hydroxy group of the zeatin side chain. The graphic shows the 6 different zeatin glucoside isomers which are identical in mass and have to be separated by chromatography for distinction.

We purchased standards for all cytokinin isomers except for cZ7G and optimized their separation by HPLC. The complete separation of all zeatin glucoside isomers is challenging. On the HSST3 column only cZ9G is well separated as a single peak, while tZ7G/tZOG and cZOG/tZ9G are co-eluting (Figure 5A). In contrast, the Waters Atlantis column yielded three clearly separated peaks for three of the five tested zeatin glucosides: cZOG, tZ9G and cZ9G, while tZ7G/tZOG are co-eluting in a fourth peak (Figure 5B).

**Figure 5:**
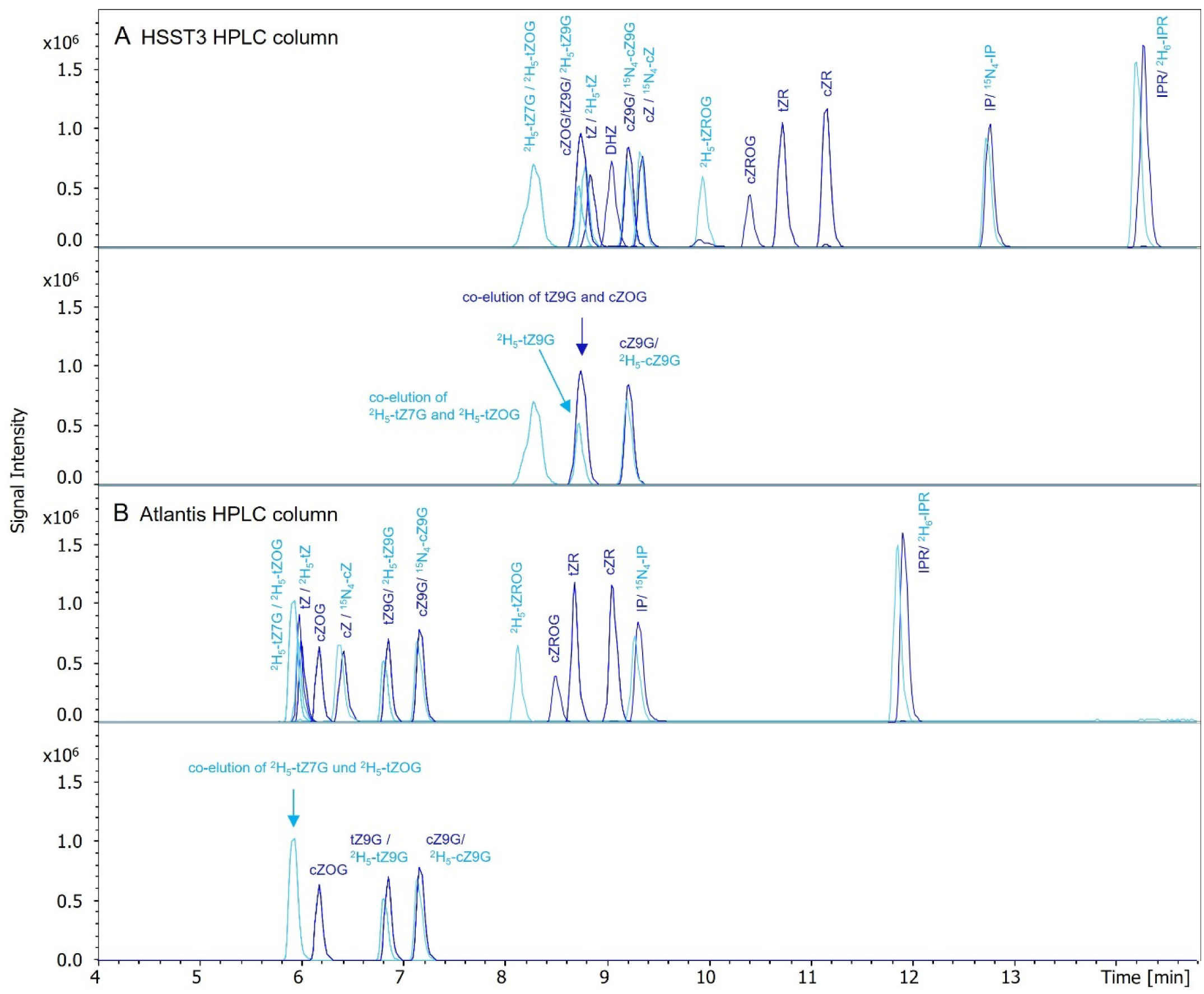
Chromatographic separation of cytokinin standards on two different HPLC columns. The analyzed mixture contained all cytokinin standards as listed in Table 2, separated on a Waters HSST3 HPLC column (A) or a Waters Atlantis HPLC column (B) using the same gradient, and measured by Q-TOF MS on a maXis 4G instrument in positive ionization mode. The lower panels of A and B show a selection of only the labeled and unlabeled zeatin-glucoside isomers. Dark blue colors represent the EICs of the respective [M+H]^+^ unlabeled standards, light blue colors represent the respective isotope-labeled standards.

Thus, when addressing biological questions where zeatin glucosides are of special interest, the separation on the Atlantis column is superior to the HSST3 column.

One additional example for *cis*/*trans* isomerism is the presence of *trans*,*trans*-ABA in addition to *cis*,*trans*-ABA which we have observed in many plant species such as Arabidopsis, rice and barley. While looking for *cis*,*trans*-ABA we sometimes detected an additional peak with the same m/z and a slightly lower RT (14.1 min vs. 14.6 min), when being separated on the HSST3 column and detected in negative ionization mode (Figure 3A and Table 1). This peak was especially prominent in rice samples. We purchased additional ABA standards and identified this peak as *trans*,*trans*-ABA. Unlike in rice or barley, *trans*,*trans*-ABA was almost at the limit of detection in unstressed Arabidopsis samples. Only during drought stress when *cis*,*trans*-ABA levels are also increased we could detect clear peaks for *trans*,*trans*-ABA. It is important to keep in mind that some phytohormone isomers may be more abundant in some plant species than others.

In addition to stereoisomers there are other phytohormones, which differ in their molecular structure but still have the same molecular mass as other related phytohormones. Examples for these are the isobaric gibberellins GA4 and GA20, or GA1 and GA34. While these are very clearly separated by LC (Figure 3A), it is important to be aware of the variety of isobaric phytohormones.

Among the conjugated auxins there are also isobaric phytohormones which share the same molecular formula, namely IAA/I3CAMe, IPA/IAAMe and IBA/IAAEt (Table 1). These phytohormones are also well separated on the HSST3 column, as can be seen in Figure 3. We emphasize the importance of an accurate assignment of the peaks detected in biological samples by comparison with the respective phytohormone standards. It is also very helpful to be aware of possible isomers or other isobaric compounds which might be present in plant extracts.

### Miscellaneous Observations during LC-Separation

#### Plant Species-specific Matrix Effects

When measuring phytohormones in different plant extracts we encountered challenges caused by the respective plant matrix. For example, 9,10-DHJA, which has been described as an internal standard for the quantification of jasmonates in Arabidopsis extracts [9], proved to be unsuitable as an internal standard for the analysis of phytohormones in barley extracts. In contrast to Arabidopsis extracts, barley extracts contained an unidentified metabolite of a highly similar m/z which co-eluted with 9,10-DHJA, thereby contaminating the internal standard peak and rendering it unsuitable for quantification. In this specific case we recognized the problem by a strong increase in peak intensity of what was allegedly 9,10-DHJA, in comparison with blank samples containing only the internal standard mix. This problem could be overcome in barley by choosing the HSST3 HPLC column where a clear separation of 9,10-DHJA and the isobaric barley-specific peak confirmed our suspicions. Alternatively, the (much more expensive) deuterated ^2^H_6_-JA could be chosen as internal standard. It is important to be aware of such problems, which can arise when dealing with non-model plant species.

#### Elution of Salicylic Acid

Some phytohormones may present unexpected challenges when their chromatographic separation is first established. In our hands one example was SA. While giving a good signal and a precise peak when separated on the HSST3 column, SA presented some difficulties when the Atlantis column was used. Here we observed strong retention time shifts over time, increasing over time, while the other phytohormones’ retention times remained stable. The retention time increased to such an extent that SA no longer eluted from the column at all after several injections, making it impossible to quantify SA in plant extracts separated on the Atlantis column with our chosen HPLC gradient. This finding was surprising to us as the Atlantis column is specifically recommended for the separation of polar acidic metabolites and this information may be of some value for other groups encountering similar problems. This problem could be fully overcome by choosing the HSST3 column for SA analysis.

### Phytohormone detection by High-Resolution Q-TOF MS versus Multiple Reaction Monitoring

Phytohormones are a chemically very diverse group of metabolites. Routinely these analytes are quantified in plant extracts by LC-MS using MRM on triple quad (QQQ) instruments. MRMs are often praised for a higher sensitivity as opposed to Q-TOF instruments, while the latter have a higher resolution and mass accuracy.

There is a large number of MRMs described in literature for known phytohormones analyzed on different triple quad instruments [e.g. 13,14]. However, MRM optimization, including the choice of transitions and voltages for parameters such as collision energy (CE) and declustering potential (DP) has to be done for each instrument individually. This requires the availability of reference standards.

This is a dilemma when faced with the task of a comprehensive phytohormone profiling of non-model plant species. Some phytohormone classes, e.g. the gibberellins or conjugated auxins comprise a very large number of molecules which can potentially be found in a plant extract. We believe that Q-TOF analysis of phytohormones is a valuable addition to MRM-based analysis on triple quads, especially when investigating non-model plant species. The high mass accuracy of a Q-TOF instrument allows the detection of phytohormones in plant extracts in MS full scan mode. Thus, to achieve a phytohormone profiling there is no restriction to compounds with known MRMs. We compared both methods back-to-back to get a detailed overview on the performance of both methods for different phytohormones.

#### High-Resolution Q-TOF MS

We analyzed phytohormones in full scan mode on a high-resolution Q-TOF mass spectrometer maXis 4G (Bruker Daltonics, Germany). The maXis 4G was improved by purchasing an upgrade leading to higher sensitivity of small molecules and to an improved resolution of 90K at 922 m/z. Data were acquired in the mass range of 100-1000 m/z.

We optimized the parameters for the ion source and the ion transfer for molecules below 600 Da. Q-TOF instruments sometimes can discriminate small ions. To increase the signal intensity of phytohormones with a very low mass, such as e.g. SA and IAA we tested which parameters have the greatest impact. On the maXis 4G Q-TOF instrument collision Rf voltage proved to be the most important parameter which had to be adjusted. A low collision RF of 300 Vpp (Vpp = peak-to-peak voltage) improves the ion transmission of small molecules and thus increases the signal intensity of small ions.

Phytohormone mass range includes small acids such a salicylic acid (m/z = 137.024) as well as larger cytokinin molecules which are conjugated to several sugar moieties (e.g. ZROG, m/z = 514.214). To achieve simultaneous analysis of both small and larger ions with optimized sensitivity we chose the manufacturers “stepping mode” employing collision Rf voltages between 300 and 500 Vpp.

Phytohormones were identified according to their m/z, with a mass tolerance of ± 0.005 mDa. Retention time was compared with isotope-labeled internal standards and/or standards injected separately, e.g. as part of quality controls or calibration curves. Due to the high mass accuracy and comparison of retention times the identification of phytohormones could be achieved at a high confidence level.

All phytohormone standards were first measured in positive and negative ionization mode. Comparison of signal intensities and signal-to-noise ratios resulted in the following ionization modes for the different phytohormone classes: Cytokinins and MeJA were analyzed in positive ion mode; SA, ABA, JA, OPDA and gibberellins were analyzed in negative ion mode. IAA was measured in both negative and positive ion mode but showed a better signal-to-noise ratio in positive ion mode, which was therefore used for IAA quantification. Most of the conjugated IAA derivatives were detectable in both negative and positive ionization mode, though all showed higher signal intensities in positive ionization mode. Amino acid conjugates of IAA showed relatively small differences in ionization efficiency between positive and negative mode. Free acids such as IBA and IPA as well as the amides IPAM and IAM clearly showed a better ionization efficiency in positive mode. IAA esters such as methylated and ethylated IAA were only detectable in positive ionization mode.

#### Multiple Reaction Monitoring

The multiple reaction monitoring (MRM) analysis was performed on a Qtrap 5500 instrument (Sciex, USA), coupled to an Agilent 1260 HPLC, using the same HPLC column and gradient as for Q-TOF MS experiments. Transitions were pre-selected based on literature and our MS2 analysis with the QTOF system and final choices were made based on signal intensity and compound specificity.

We measured phytohormone standards on the maXis 4G Q-TOF mass spectrometer in MS2 mode to obtain fragment spectra. These fragment spectra show all fragments of the compound. The same standards were then measured in direct infusion mode on the Sciex Qtrap 5500 instrument, conducting the compound optimization procedure. Thus instrument-specific parameters were optimized for up to four most suitable transitions. One parameter which influences signal intensity is the declustering potential (DP). DP was optimized to obtain a high analyte signal while avoiding premature fragmentation of the analyte in the source (“in-source fragmentation”). The declustering potential was optimal for all phytohormones over a relatively broad range and was therefore kept very similar between -60 V to -50 V for all MRMs in negative mode and at +50 V in positive mode. Adapting the cell entrance potential (CEP) and cell exit potential (CXP) had neglectable effects, thus these were kept constant at +10V/-10V (CEP) and +15V/-15V (CXP).

The parameter which had strongest impact on signal intensities of individual MRMs was the collision energy (CE). CEs have to be assessed separately for each MRM for all phytohormones. Collision energies were ramped for up to four candidate MRMs during the manufacturer’s compound optimization procedure. If possible two transitions were chosen which showed the highest intensities at their optimized collision energy. Transitions and optimized collision energies are shown in Table 3. An additional important aspect to consider was selectivity: when several transitions were suitable, those which were specific for a given phytohormone were preferred. This decision was based also on the information of the MS2 spectra from the QTOF system. This is especially helpful to distinguish isobaric phytohormones. For the same reason two transitions were selected where possible to improve selectivity. Quantification was done on the sum of two MRMs where applicable.

There are several publications which list transitions and instrument parameters of the phytohormones analyzed in their studies. However, it is important to know that parameters have to be adjusted individually to the used instruments to obtain optimal results. Furthermore, it can be difficult to obtain information on suitable MRM transitions for isotopically labeled standards from the literature. The availability of isotopically labeled standards is increasing; therefore, some standards may not have been used in previous literature. It is of great importance when choosing the MRMs to be aware of the position of the isotope label on the molecule. Literature giving supplemental information on the fragmentation patterns of different labeled phytohormones in comparison to their unlabeled counterparts is very valuable [13]. Table 3 shows the two most suitable MRMs selected for unlabeled and isotope-labeled phytohormone standards used in our method as well as the corresponding CEs in positive or negative ionization mode. Due to the ability of QQQ instruments for fast polarity switching, both positive and negative ionization experiments could be performed within one single acquisition run.

**Table 3:** Optimized transitions of phytohormones for multiple reaction monitoring (MRM) on the Sciex Qtrap 5500. In each line the phytohormone of interest, the preferred ESI mode and the MRM transitions used for quantification with corresponding collision energies (CE) are shown. Where applicable the respective isotope-labeled internal (I.S.) and the corresponding MRM transitions and collision energies are shown in columns 5-7.

| Standard | ESI mode | MRM transitions | CE1<br>CE2 | Internal standard (I.S.) | MRM transitions | CE1<br>CE2 |
| --- | --- | --- | --- | --- | --- | --- |
| ABA |  |  |  |  |  |  |
| (±)-*cis,trans*-Absciscic acid (ABA) | - | 263.1 > 153.1<br>263.1 > 219.2 | -15<br>-18 | <sup>2</sup>H<sub>6</sub> (+)-*cis,trans*-Absciscic acid (<sup>2</sup>H<sub>6</sub>-ABA) | 269.1 > 159.1<br>269.1 > 207.1 | -15<br>-22 |
| (+)-*trans,trans*-Absciscic acid (S-tt-ABA) | - | 263.1 > 153.1<br>263.1 > 219.2 | -15<br>-18 |  |  |  |
| (±)-*cis,trans*-Absciscic acid glucosyl ester (ABA-Glc) | + | 427.3 > 265.1<br>427.3 > 247.1 | 9<br>18 |  |  |  |
| Gibberellins |  |  |  |  |  |  |
| Gibberellin A1 (GA1) | - | 347.1 > 259.0<br>347.1 > 303.0 | -25<br>-25 | <sup>2</sup>H<sub>4</sub>-Gibberellin A1 (<sup>2</sup>H<sub>4</sub>-GA1) | 351.2 > 263.0<br>351.2 > 307.0 | -28<br>-28 |
| Gibberellic acid (GA; GA3) | - | 345.1 > 239.0<br>345.1 > 143.0 | -19<br>-34 | <sup>2</sup>H<sub>4</sub>-Gibberellin A1 (<sup>2</sup>H<sub>4</sub>-GA1) |  |  |
| Gibberellin A4 (GA4) | - | 331.2 > 243.0<br>331.2 > 287.0 | -24<br>-25 | <sup>2</sup>H<sub>2</sub>-Gibberellin A4 (<sup>2</sup>H<sub>2</sub>-GA4) | 333.2 > 245.2<br>333.2 > 289.2 | -26<br>-26 |
| Gibberellin A8 (GA8) | - | 363.1 > 275.0<br>363.1 > 257.0 | -24<br>-23 |  |  |  |
| Gibberellin A9 (GA9) | - | 315.2 > 271.0<br>315.2 > 253.0 | -27<br>-34 | <sup>2</sup>H<sub>2</sub>-Gibberellin A9 (<sup>2</sup>H<sub>2</sub>-GA9) | 317.2 > 273.0<br>317.2 > 255.0 | -25<br>-35 |
| Gibberellin A19 (GA19) | - | 361.2 > 317.0<br>361.2 > 273.0 | -33<br>-36 | <sup>2</sup>H<sub>2</sub>-Gibberellin A19 (<sup>2</sup>H<sub>2</sub>-GA19) | 363.2 > 319.0<br>363.2 > 275.0 | -34<br>-38 |
| Gibberellin A20 (GA20) | - | 331.2 > 243.0<br>331.2 > 287.0 | -24<br>-25 | <sup>2</sup>H<sub>2</sub>-Gibberellin A20 (<sup>2</sup>H<sub>2</sub>-GA20) | 333.2 > 245.2<br>333.2 > 289.2 | -26<br>-26 |
| Gibberellin A34 (GA34) | - | 347.2 > 241.0<br>347.2 > 259.0 | -26<br>-23 | <sup>2</sup>H<sub>2</sub>-Gibberellin A34 (<sup>2</sup>H<sub>2</sub>-GA34) | 349.2 > 243.0<br>349.2 > 261.0 | -25<br>-25 |
| Gibberellin A44 (GA44) | - | 345.2 > 301.0<br>345.2 > 273.0 | -32<br>-33 | <sup>2</sup>H<sub>2</sub>-Gibberellin A44 (<sup>2</sup>H<sub>2</sub>-GA44) | 347.2 > 303.0<br>347.2 > 275.0 | -35<br>-35 |
| Gibberellin A53<sup>a</sup> (GA53) | - | 347.2 > 303.0<br>347.2 > 187.9 | -37<br>-48 | <sup>2</sup>H<sub>2</sub>-Gibberellin A53 (<sup>2</sup>H<sub>2</sub>-GA53) | 349.2 > 304.9<br>349.2 > 189.0 | -37<br>-48 |
| Salicylic Acid and Others |  |  |  |  |  |  |
| Salicylic acid(SA) | - | 137.0 > 92.8<br>137.0 > 64.9 | -20<br>-37 | <sup>2</sup>H<sub>4</sub>-Salicylic acid (<sup>2</sup>H<sub>4</sub>-SA) | 141.1 > 97.1<br>141.1 > 69.1 | -22<br>-40 |
| Cinnamic acid (CA) | - | 147.0 > 103.1 | -15 |  |  |  |
| Jasmonates |  |  |  |  |  |  |
| (±)-Jasmonic acid (JA) | - | 209.1 > 59.0 | -15 | <sup>2</sup>H<sub>6</sub>-(±)-Jasmonic acid (<sup>2</sup>H<sub>6</sub>-JA) | 215.2 > 59.0 | -15 |
|  | - |  |  | (±)-9,10-Dihydro-jasmonic acid (9,10-DHJA) | 211.1 > 59.0 | -15 |
| (±)-Jasmonic acid methyl ester (MeJA) | + | 225.1 > 151.1<br>225.1 > 133.1 | 16<br>18 |  |  |  |
| N-[(−)-Jasmonoyl]-(S/L)-isoleucine<sup>b</sup> (Jaile) | - | 322.2 > 130.0<br>322.2 > 172.1 | -27<br>-22 | <sup>2</sup>H<sub>6</sub>-N-[(−)-Jasmonoyl]-(L)-isoleucine (<sup>2</sup>H<sub>6</sub>-Jaile) | 328.2 > 130.0<br>328.2 > 172.1 | -27<br>-22 |
| N-[(−)-Jasmonoyl]-(S/L)-isoleucine<sup>b</sup> (Jaile) | + | 324.0 > 151.1<br>324.0 > 278.0 | 22<br>17 | <sup>2</sup>H<sub>6</sub>-N-[(−)-Jasmonoyl]-(L)-isoleucine (<sup>2</sup>H<sub>6</sub>-Jaile) | 330.1 > 157.1<br>330.1 > 284.0 | 24<br>19 |
| cis-12-oxo-Phytodienoic acid (cis-OPDA) | - | 291.2 > 165.1<br>291.2 > 247.0 | -25<br>-24 | <sup>2</sup>H<sub>5</sub>-*cis*-12-oxo-Phyto-dienoic acid (<sup>2</sup>H<sub>5</sub>-OPDA) | 296.2 > 170.0 | -25 |

| Auxins |  |  |  |  |  |  |
| --- | --- | --- | --- | --- | --- | --- |
| Indole-3-acetic acid (IAA) | + | 176.1 > 130.1<br>176.1 > 103.1 | 20<br>43 | <sup>2</sup>H<sub>5</sub>-Indole3-acetic acid (<sup>2</sup>H<sub>5</sub>-IAA) | 181.1 > 134.0<br>181.1 > 106.0 | 20<br>43 |
| Indole-3-acetyl-aspartic acid<sup>b</sup> (IAAsp) | + | 291.1 > 130.1<br>291.1 > 134.0 | 30<br>16 | <sup>2</sup>H<sub>5</sub>-Indole-3-acetyl-<sup>15</sup>N<sub>4</sub>-aspartic acid (<sup>2</sup>H<sub>5</sub><sup>15</sup>N<sub>4</sub>-IAAsp) | 297.1 > 134.0 | 30 |
| Indole-3-acetyl-aspartic acid<sup>b</sup> (IAAsp) | - | 289.0 > 131.8<br>289.0 > 173.8 | 23<br>20 | <sup>2</sup>H<sub>5</sub>-Indole-3-acetyl-<sup>15</sup>N<sub>4</sub>-aspartic acid (<sup>2</sup>H<sub>5</sub><sup>15</sup>N<sub>4</sub>-IAAsp) | 295.0 > 132.8<br>295.0 > 178.9 | -23<br>-21 |
| Indole-3-acetyl-glutamic acid (IAGlu) | + | 305.1 > 130.1<br>305.1 > 148.1 | 28<br>18 |  |  |  |
| Indole-3-acetyl-alanine (IAAla) | + | 247.1 > 130.1<br>247.1 > 90.1 | 30<br>15 |  |  |  |
| Indole-3-acetyl-valine (IAVal) | + | 275.1 > 130.1<br>275.1 > 229.0 | 32<br>16 |  |  |  |
| Indole-3-acetic acid methyl ester (IAAMe) | + | 190.1 > 130.1<br>190.1 > 77.0 | 22<br>60 |  |  |  |
| Indole-3-acetyl-tryptophane (IATrp) | + | 362.1 > 130.1<br>362.1 > 205.0 | 40<br>18 |  |  |  |
| Indole-3-acetyl-isoleucine (IAIleu) | + | 289.2 > 130.1<br>289.2 > 243.0 | 36<br>16 |  |  |  |
| Indole-3-acetyl-phenylalanine (IAPhe) | + | 323.1 > 130.1<br>323.1 > 166.0 | 30<br>18 |  |  |  |
| Indole-3-acetic acid ethyl ester (IAAEt) | + | 204.1 > 130.1<br>204.1 > 77.0 | 27<br>63 |  |  |  |
| Indole-3-acetamide (IAM) | + | 175.1 > 130.1<br>175.1 > 158.0 | 22<br>12 |  |  |  |
| Indole-3-propionamide (IPAM) | + | 189.1 > 130.1<br>189.1 > 172.0 | 23<br>12 |  |  |  |
| Indole-3-carboxylic acid (I3CA) | + | 162.1 > 116.0<br>162.1 > 91.0 | 29<br>36 |  |  |  |
| Indole-3-carboxylic acid methyl ester (I3CAME) | + | 176.1 > 144.0<br>176.1 > 116.0 | 18<br>33 |  |  |  |
| Indole-3-acetonitrile (I3ACN) | + | 157.1 > 130.1<br>157.1 > 117.1 | 15<br>30 | <sup>2</sup>H<sub>4</sub>-Indole-3-acetonitrile (I3ACN) | 157.1 > 133.1<br>157.1 > 121.1 | 14<br>29 |
| Indole-3-propionic acid (IPA) | + | 190.1 > 130.1<br>190.1 > 76.9 | 26<br>62 |  |  |  |
| Indole-3-butyric acid (IBA) | + | 204.1 > 130.1<br>204.1 > 186.1 | 26<br>26 |  |  |  |
| Cytokinins |  |  |  |  |  |  |
| *trans*-Zeatin (tZ) | + | 220.1 > 136.1<br>220.1 > 119.0 | 26<br>46 | <sup>2</sup>H<sub>5</sub>-*trans*-Zeatin (<sup>2</sup>H<sub>5</sub>-tZ) | 225.2 > 137.1<br>225.2 > 149.1 | 25<br>24 |
| *cis*-Zeatin (cZ) | + | 220.1 > 136.1<br>220.1 > 119.0 | 26<br>46 | <sup>15</sup>N<sub>4</sub>-*cis*-Zeatin (<sup>15</sup>N<sub>4</sub>-cZ) | 224.1 > 140.1<br>224.1 > 122.0 | 24<br>46 |
| Dihydrozeatin (DHZ) | + | 222.1 > 136.1<br>220.1 > 148.1 | 29<br>29 |  |  |  |
| *trans*-Zeatin-O-glucoside (tZOG) | + | 382.2 > 220.1<br>382.2 > 136.1 | 24<br>40 | <sup>2</sup>H<sub>5</sub>-*trans*-Zeatin-O-glucoside (<sup>2</sup>H<sub>5</sub>-tZOG) | 387.2 > 225.1<br>387.2 > 137.1 | 27<br>40 |
| *trans*-Zeatin-7-glucoside<sup>a</sup> (tZ7G) | + |  |  | <sup>2</sup>H<sub>5</sub>-*trans*-Zeatin-7-glucoside (<sup>2</sup>H<sub>5</sub>-tZ7G) | 387.2 > 225.1<br>387.2 > 137.1 | 27<br>40 |
| *cis*-Zeatin-O-glucoside (cZOG) | + | 382.2 > 220.1<br>382.2 > 136.1 | 24<br>40 |  |  |  |
| *trans*-Zeatin-9-glucoside (tZ9G) | + | 382.2 > 220.1<br>382.2 > 136.1 | 24<br>40 | <sup>2</sup>H<sub>5</sub>-*trans*-Zeatin-9-glucoside (<sup>2</sup>H<sub>5</sub>-tZ9G) | 387.2 > 225.1<br>387.2 > 137.1 | 27<br>40 |
| *cis*-Zeatin-9-glucoside (cZ9G) | + | 382.2 > 220.1<br>382.2 > 136.1 | 24<br>40 | <sup>15</sup>N<sub>4</sub>-*cis*-Zeatin-9-glucoside (<sup>15</sup>N<sub>4</sub>-cZ9G) | 386.2 > 224.1<br>386.2 > 140.0 | 24<br>40 |
| *trans*-Zeatin-O-glucoside riboside (tZROG) | + | 514.2 > 382.2<br>514.2 > 220.1 | 26<br>38 | <sup>2</sup>H<sub>5</sub>-*trans*-Zeatin-O-glucoside riboside (<sup>2</sup>H<sub>5</sub>-tZROG) | 519.2 > 387.2<br>519.2 > 225.1 | 26<br>38 |
| *cis*-Zeatin-O-glucoside riboside (cZROG) | + | 514.2 > 382.2<br>514.2 > 220.1 | 26<br>38 |  |  |  |
| *trans*-Zeatin riboside (tZR) | + | 352.2 > 220.1<br>352.2 > 136.1 | 24<br>40 |  |  |  |
| *cis*-Zeatin riboside (cZR) | + | 352.2 > 220.1<br>352.2 > 136.1 | 24<br>40 |  |  |  |
| N6-Isopentenyl-adenine (IP) | + | 204.1 > 136.1<br>204.1 > 148.1 | 22<br>16 | <sup>15</sup>N<sub>4</sub>-N6-Isopentenyl-adenine (<sup>15</sup>N<sub>4</sub>-IP) | 208.1 > 140.0<br>208.1 > 152.0 | 22<br>18 |
| N6-Isopentenyl-adenosine (IPR) | + | 336.2 > 204.1<br>336.2 > 136.0 | 24<br>40 | <sup>2</sup>H<sub>6</sub>-N6-Isopentenyl-adenosine (<sup>2</sup>H<sub>6</sub>-IPR) | 342.2 > 210.0<br>342.2 > 137.0 | 24<br>37 |

ESI= electrospray ionization; CE= collision energy; I.S. = internal standard; <sup>2</sup>H<sub>x</sub>= deuterated with x deuterium atoms
<sup>a</sup> for GA53 and tZ7G only isotope-labeled phytohormone standards were available, therefore transitions were optimized with the isotope-labeled equivalent and the thus obtained optimized energies are displayed in this table.
<sup>b</sup> for Jaile and IAA<sub>sp</sub>, MRM transitions for both positive and negative ESI mode are displayed because both ESI modes can be employed with similar efficiency and fragmentation behavior is strongly affected by the ESI mode.
<sup>c</sup> where no I.S. is depicted, no isotope-labeled I.S. were available and related I.S. were used for quantification.

#### High-Resolution Q-TOF MS versus Multiple Reaction Monitoring

Both strategies, MRM as well as HRMS using Q-TOFs, have distinct advantages for the detection and identification of phytohormones. Q-TOF MS is a highly efficient tool to obtain a comprehensive overview on the phytohormone status of a plant without extensive prior knowledge and detection method optimization. Thus, Q-TOF MS is our method of choice when a novel plant species or unusual plant tissues are investigated. Information obtained in these analyses can then be used to build a tailored sensitive MRM method to routinely analyze phytohormones in these samples.

The sensitivity of both methods is analyte-dependent. Therefore, it has to be determined for all compounds individually. Sensitivity of the MRM method can be superior for phytohormones where specific transitions with good signal intensities are available. Examples are JA, JA-Ileu and GA3 in negative mode and IAA and zeatins in positive mode. In our hands, Q-TOF MS is equally sensitive for some phytohormones such as *trans*, *trans*-ABA and some gibberellins (GA44, GA20, GA4).

The differences in sensitivity between GA3 and other gibberellins like GA44, GA20 and GA4 may be unexpected and arise from a different fragmentation behavior upon collision induced dissociation (CID). While GA3 readily yields several high intensity fragments which can be used to establish a sensitive MRM method, the GA4 precursor does not fragment easily upon lower collision energies (CEs) and signal intensity is lost upon higher CEs, resulting in low sensitivity of the MRM method.

The selectivity of MRMs proves an advantage for some phytohormones when high backgrounds are encountered in plant extracts. We observed this for example in the analysis of JA and OPDA in barley meristem, where MRMs were advantageous due to reduced background.

However, selectivity can be compromised when related phytohormones yield similar fragment ions. This is the case for IAA and its conjugates. All of them yield m/z = 130 as the major fragment ion in positive mode. Thus, using a transition from a parent ion to the product ion m/z = 130 is a suitable transition to quantify auxins containing an IAA backbone, and this general transition is suitable to screen for members of this compound class. However, such a transition is not helpful to differentiate between different members of this class. For example, IPA and IAAMe both have the same parent ion and upon CID both yield m/z = 130 as the major product ion.

Similarly, zeatins consistently produce the same fragments, which makes is easy to screen for phytohormones containing the zeatin backbone, but their profiling requires careful comparison of retention times to distinguish between isomers. Here, an important pitfall to avoid is possible in-source decay of glucosylated zeatins, which results in the loss of glucose and yields zeatin fragments in the ion source. These in turn produce product ions upon CID that cannot be distinguished from those of free zeatin. Using Q-TOF MS such an in-source decay would be easily detected by inspecting the full scan mass spectra, showing the presence of the corresponding zeatin-glucoside ion. However, when an MRM method is employed that does not specifically target zeatin-glucosides, these may easily be confused with free zeatin, especially when the retention times during HPLC separation are very similar.

Another advantage of Q-TOF MS is the possibility to re-evaluate data after the measurements have been completed. If biological questions arise which warrant the analysis of additional phytohormones or other metabolites of interest, it is possible to search for specific m/z in previously measured samples and evaluate these data. This is possible because all data are acquired in an unbiased manner. This makes Q-TOF MS preferable for plant species or tissues which are not routinely analyzed and where unexpected phytohormones may be encountered. In conclusion, Q-TOF MS allows for unbiased phytohormone profiling due to no limitations by MRMs. Thus Q-TOF MS is a valuable method to support MRM-based routine analysis.

### Additional Information for Phytohormone Profiling and Quantification

The correct identification of phytohormones is the prerequisite for all quantitative analyses that follow. There are three major characteristic properties that can be used for compound identification of metabolites using LC-MS analysis:

- Retention Time (HPLC)
- MS1: Accurate mass (Q-TOF)/mass of precursor ion (QQQ)
- MS2: Fragmentation spectra (Q-TOF) /mass of product ion (QQQ)

Retention times are determined by comparison of plant extracts with commercially available phytohormone standards. These have to be purchased, can be expensive and are not always available for all phytohormone species. The analysis of a single phytohormone standard will provide a retention time for the chosen solvent gradient on a specific HPLC column. Retention times for all phytohormone standards analyzed in this study can be found in Table 1.

It is important that standards are analyzed separately from other standards during method establishment to rule out a possible confusion with other standards. This is especially important for phytohormones because of the large number of isomers or other isobaric compounds. In this study alone, we have analyzed a range of phytohormone standards which have at least one isobaric counterpart:

- GA20 and GA4
- GA1 and GA34
- IPA and IAAMe
- IAA and I3CAMe
- IBA and IAAEt
- *trans*,*trans*-ABA and *cis,trans*-ABA
- *trans*-zeatin and *cis*-zeatin
- tZOG and cZOG, tZ9G and cZ9G

Once the retention times have been established for each standard separately, they can be measured as standard mixes during subsequent analyses, e.g. as part of quality controls (QCs) or calibration curves. Matrix effects such as pH-shifts or interactions between metabolites may affect the retention time of phytohormones in a plant extract. Also, the solvent in which the phytohormones are injected can have an effect on the retention time. These effects have to be considered when comparing phytohormone standards injected separately with phytohormones in a plant extract. When in doubt, spiking the standard into a plant extract can give valuable information on the effect of the plant extract matrix on the retention time of a given phytohormone.

When triple quads (QQQ) are used, their mass resolving power allows the distinction of phytohormones that differ in mass by approx. 1 Da. This is not sufficient to confidently identify analytes based on the mass of the precursor alone. Therefore, analysis is usually done with MRMs, requiring information on the mass of the product ions as well.

In contrast, HRMS with a mass resolution of ≥ 60.000 can provide a high confidence in assigning the elemental composition of an analyte. This allows the distinction of phytohormones with different sum formulas at MS1 level, which is a great advantage over triple quad analysis. Additional acquisition of MS2 data using HRMS is also possible and can support the identification of known phytohormones and even enable the structural elucidation of novel phytohormones.

#### The Importance of Blank Controls

To ensure correct quantification of phytohormones it is important that the signals obtained for a detected phytohormone truly correspond to the phytohormone in question. However, there may, rarely, be contaminants of non-plant origin which can yield a highly similar background signal in the mass spectrometer. To avoid the false quantification of these background signals it is important to analyze blank samples containing only the internal standard, which have undergone the same extraction process as the plant samples. If background signals are indeed detected in these blank samples, they may either be ignored (e.g. if peaks occur at a different retention time) or the respective peak areas can be subtracted from plant samples. We have observed both scenarios: an additional peak regularly occurred in the blank with the exact same m/z as GA19 at a different retention time, as well as a constant low signal with the same m/z and retention time as isopentenyladenine (IP). With this knowledge we could ignore the false GA19 peak and ensure that IP was only quantified in plant samples where the peak area exceeded that found in blank samples.

#### Quantification

To ensure reliable quantification of phytohormones it is important to account for extraction efficiency, stability and solubility of phytohormones and matrix effects in different plant extracts. In addition to using an optimized extraction protocol for phytohormone analysis to minimize losses, isotope-labeled internal standards (I.S.) were used to identify effects on signal intensities due to phytohormone losses during extraction or due to matrix effects. The heterogenic chemical nature of the large number of phytohormones means that these effects may affect some phytohormones to a different extent than others. Thus, the use of as many isotope-labeled standards as possible increases the reliability of the analyses. We have purchased a large number of isotope-labeled standards (Table 1/2 and Table 3) to use for phytohormone quantification at a concentration of 75 nM in the final sample. For some phytohormones no corresponding isotopically labeled standard was available. To overcome this problem structurally related standards were used as an alternative, for example ^2^H2-GA19 was used for GA8 and ^2^H5-tZOG for cZOG.

For absolute quantification, the same internal standard mix were used at a fixed concentration of 75 nM, together with unlabeled phytohormone standards used at concentrations ranging from 0.1 nM to 750 nM, to produce calibration curves, which were determined for each phytohormone separately. Peak areas of unlabeled and labeled phytohormones were determined with the softwares QuantAnalysis (version 5.3, Bruker Daltonics) or Analyst (version 1.7.3, Sciex) and further calculations were done in Microsoft Excel 2010. Area ratios were calculated by dividing the peak area of an unlabeled phytohormone by the peak area of its corresponding stable-isotope labeled standards (see Table 1). Absolute quantification of phytohormones was conducted based on the slopes of calibration curves. Only in the case of GA53, where the isotope-labeled internal standard was available, but the unlabelled phytohormone standard was not, the slope of a calibration curve based on area ratios from GA19 relative to the I.S. ^2^H_2_-GA53 was used for absolute quantification. To make data comparable with existing literature, we calculated final phytohormone concentrations in nmol/g fresh weight and ng/g fresh weight.

## Summary

The method we described is not specialized to one phytohormone class, but rather provides the means to do a comprehensive screening for as many phytohormones as possible. We compared LLE versus SPE-separation, optimized separation of phytohormone classes using different HPLC columns and compared Q-TOF full scan MS versus MRM as detection methods. We also shared detailed information of observations and pitfalls that can be encountered using these methods.

Our proposed optimized method consists of an optimized liquid-liquid extraction protocol which yields a very good recovery of the vast majority of the 50 tested phytohormones. This method is suitable for the extraction from small tissue samples of 5-50 mg plant material and can be used for the parallel extraction of large numbers of samples. Our optimized HPLC separation employs a C18 XSelect™ HSST3 column (Waters) for a comprehensive screening of phytohormones, and a C18 Atlantis™ Premier BEH column (Waters) when special focus is laid on the separation of zeatin-glucoside isomers. We provide detailed information on the presence and separation of phytohormone isomers and isobars among all phytohormone classes including retention times. Regarding the detection method, we share our findings concerning the optimal ionization parameters for all investigated phytohormones, for ionization in positive as well as negative mode. We describe optimized MRM parameters including transitions and CEs for all analyzed phytohormones.

We found great value in Q-TOF MS analyses for unbiased phytohormone profiling compared to targeted transitions limitating the scope of identification by MRMs. In our hands, Q-TOF MS is the method of choice to investigate phytohormones in non-model plant species. Information obtained from Q-TOF experiments can then subsequently be used to build a MRM-based routine analysis.

## Limitations

We have described our optimized method for the comprehensive and reliable analysis of phytohormones comprising the classes of auxins, cytokinins, GAs, ABA, SA and jasmonates. Due to the highly diverse physicochemical properties of the phytohormones this method is a compromise and not optimized to one specific phytohormone. Some bioactive phytohormones are extremely low abundant and can be below the limit of detection. In such cases, if very specific phytohormones from only one class are of interest, another protocol may be established with method details tailored specifically to the physicochemical properties of the phytohormone of interest.

## Ethics statements

Not applicable

## CRediT author statement

**Vera Wewer:** Conceptualization, Methodology, Investigation, Validation, Writing-Reviewing and Editing

**Nadine Dyballa-Rukes:** Investigation, Reviewing and Editing

**Sabine Metzger:** Supervision, Conceptualization, Methodology, Reviewing and Editing

## Acknowledgments

We would like to thank Dr. Meike Siebers (Institute for Plant Genetics, Heinrich-Heine-University Düsseldorf) and Daniela Krüger (CEPLAS Plant Metabolism and Metabolomics Facility, University of Cologne) for their help during initial phytohormone extraction and purification experiments.

Figures 1, 2 and the graphical abstract were created with the help of BioRender.com.

This work was funded by the Deutsche Forschungsgemeinschaft (DFG, German Research Foundation) under Germanýs Excellence Strategy – EXC-2048/1 – project ID 390686111

